# Bayesian optimization of cortical neuroprosthetic vision using perceptual feedback

**DOI:** 10.1101/2025.01.24.634528

**Authors:** Burcu Küçükoğlu, Leili Soo, David Leeftink, Fabrizio Grani, Cristina Soto Sanchez, Umut Güçlü, Marcel van Gerven, Eduardo Fernandez

## Abstract

The challenge in cortical neuroprosthetic vision is determining the optimal, safe stimulation patterns for the visual cortex in order to evoke the desired perception in blind individuals—specifically, light perceptions known as phosphenes. Currently, clinical studies gain insights into the perceptual characteristics of the perceived phosphenes by asking for descriptions of provided stimulation protocols. However, the huge parameter space for multi-electrode stimulation settings makes it difficult to draw conclusions about the optimality of the stimulation patterns that lead to well-perceived phosphenes. A systematic search in the parameter space of the electrical stimulation is needed to achieve good perception. Bayesian optimization (BO) is a framework for finding optimal parameters efficiently. Using the patient’s scoring of the perception as feedback, a model of the patient’s response based on iteratively generated stimulation protocols can be built to maximize perception quality. A patient implanted with an intracortical 96-channel microelectrode array in their visual cortex was tested by iteratively presenting stimulation protocols, generated via BO for the first and random generation (RG) for the second experiment. Whereas standard BO methods do not scale well to problems with over a dozen inputs, we propose to optimize a set of 40 electrode currents using trust region-based BO. The generated protocols determine which electrodes are concurrently stimulated from the set and with how much current from a range of 0-50 *µ*A, on a maximum total current constraint of 500 *µ*A. The patient provided feedback for each stimulation based on their liking of the perception quality on a Likert scale, where a score of 7 indicated the highest quality and 0 no perception. In the BO experiment, the patient perception quality ratings gradually converged on higher values compared to the RG experiment. Similarly, gradually higher total current values were chosen by BO, in line with the observed preference of patients for higher currents due to brighter phosphenes. Finally, the electrodes that were observed to be more effective in producing phosphene perception in previous studies were gradually chosen more by BO also with the allocation of higher current values. This study demonstrates the power of BO in converging to optimal stimulation protocols based on patient feedback, providing a more efficient search for stimulation parameters for clinical studies.

## 1 Introduction

Cortical neuroprostheses offer a promising approach to artificial restoration of vision in blind people. By direct electrical stimulation of the visual cortex through an electrode implant, light percepts called phosphenes can be evoked in the patient’s visual field (Brindley and Lewin, 1968; Dobelle and Mladejovsky, 1974). The key challenge in evoking the desired perception lies in providing the optimal stimulation patterns in a safe manner. Finding optimal stimulation patterns that lead to meaningful and informative perception without over-stimulation of the cortex is of interest for both clinical and simulated phosphene vision studies.

Studies of simulated phosphene vision (Chen et al., 2009a,b) have emerged as a means to visualize the supposed perceptual effect of a proposed stimulation, ideally based on biological modeling of its consequences on the brain (van der Grinten et al., 2024; Fine and Boynton, 2024; Granley et al., 2022b). By allowing for the evaluation of a stimulation pattern’s visual outcome (de Ruyter van Steveninck et al., 2022b, 2024) simulated phosphene vision also enables optimization of stimulation parameters virtually in a controlled test bed without the limitations from ethical and safety considerations of testing on patients. Therefore, relatively high resolution patterns through stimulation of multiple electrodes can be freely tested, based on optimizations of even complex end-to-end machine learning algorithms (de Ruyter van Steveninck et al., 2022a; Granley et al., 2022a; Küçükoğlu et al., 2022). Such algorithms tend to have a high number of parameters to be optimized that require large amounts of data for training and possibly a long pre-training time. However, there is no way of knowing the exact perceived effect of stimulation on the patients unless testing with them directly. Clinical studies, on the other hand, are focusing on understanding the perceptual effects of stimulation protocols administered based on descriptions from the patients regarding the shape, color, brightness, and localization of the phosphenes evoked in single electrode stimulation settings (Fernandez et al., 2021; Brindley and Lewin, 1968; Schmidt et al., 1996; Dobelle et al., 1974). These single electrode stimulations also help identify the threshold current needed for an electrode to induce perception (Chen et al., 2020). Typically, an arbitrary multiplier to this threshold value is applied when administering current to ensure phosphene perception, albeit with no guarantee of good-quality perception, due to a mere focus on the existence of perceptual detection rather than quality. For multi-electrode settings (Bosking et al., 2022; Oswalt et al., 2021), electrodes are combined in shapes, e.g. in a line, circle, or a letter, to check for the recognition of the intended pattern (Fernandez et al., 2021), or its orientation (Chen et al., 2020). Once a satisfactory description of the intended pattern is found, the stimulation protocol that gave rise to it is recorded for future use. Another way of testing for stimulation in multi-electrode settings is random selection of electrodes to concurrently stimulate along with the random assignment of their current values under safety constraints. However, this is done minimally, as it is hard to draw conclusions about the optimality of stimulation protocols in a huge parameter space based only on verbal descriptions, unless a systematic approach is taken to a process that otherwise could be tedious.

The gap between the testing conditions of clinical and simulated phosphene vision studies demonstrates the need for a systematic approach for searching the space of parameters for multi-electrode stimulation in patients in an informed manner. This would enable clinical studies to take a step further towards testing more complex stimulation protocols that currently tend to emerge from simulated phosphene vision studies’ safe yet virtual optimization conditions.

### Our proposal

We envision an optimization setup based on direct patient feedback on stimulation protocols generated in an intelligent and autonomous manner, to search systematically through the space of parameters in a fast and efficient way given safety constraints. For this purpose, we propose Bayesian optimization as a fast and theoretically optimal method of parameter optimization in clinical settings. As an initial step towards this vision, here we present a procedure to optimize stimulation protocols involving a selection of electrodes to stimulate and their current levels, based on direct patient feedback on perceptual quality. To our knowledge, this is the first automated optimization procedure in the field of neuroprosthetic vision that is based on direct patient feedback. Such a systematic search procedure for optimal stimulation protocols could make early experimentation with a patient implanted more efficient and additionally informative on perception quality, especially for multi-electrode stimulation settings.

### Bayesian optimization

Bayesian optimization (BO) (Shahriari et al., 2016) is an approach for solving black-box functions that converges quickly to globally optimal parameters with as few function evaluations as possible by searching within constraints given. This makes BO ideal for safety critical settings like neurostimulation optimization where evaluating the objective function is also expensive. It works by iteratively presenting data points and observing its quantitative evaluation in response, in order to fit a surrogate model capturing the relationship between the inputs and outputs of the black-box function. The model is updated at each iteration after the response to the newly generated data point, based on all past evaluations. The updated surrogate model in turn helps generate the next data point to be presented, using an acquisition function that reflects the surrogate’s optimal or uncertain areas to ensure a balance between exploration and exploitation in the search space. The generated data points gradually converge to data points that maximize the evaluation as the model converges to optimal parameters.

BO has been previously used for various safety-critical contexts from robotics to neurostimulation (Sui et al., 2015; Berkenkamp et al., 2021; Cooper and Netoff, 2022). In the context of neurostimulation, the use of BO involves optimization of parameters *in vivo* for deep brain stimulation in rats (Cole et al., 2024) and in Parkinson’s disease patients (Sarikhani et al., 2022) within safety constraints, for tACS to evoke phosphenes in seeing human subjects steered by their preference ratings (Lorenz et al., 2019), and for stimulation on motor cortex (Choinière et al., 2024) in rats and monkeys (Bonizzato et al., 2023). BO has also been utilized for *in silico* parameter optimization for deep brain stimulation (Nagrale et al., 2023; Cooper and Netoff, 2022; Grado et al., 2018), for offline recalibration of neuromodulation parameters over time in somatosensory neuroprosthesis (Aiello et al., 2023), for offline optimization of peripheral motor neuroprosthesis stimulation from rat and monkey data (Losanno et al., 2021), for modeling spinal cord stimulation preferences from patient data to improve therapeutic outcomes (Zhao et al., 2021), and finally for optimization of patient-specific parameters in retinal implants (Granley et al., 2023). However, these studies focused on for a maximum of 5 parameters (Bonizzato et al., 2023) for *in vivo* and 13 parameters (Granley et al., 2023) for *in silico* optimization settings.

### Our approach

In the context of neurostimulation optimization of our study, the objective function can be considered as a function of phosphene perception quality as defined by patient feedback, with its input being the stimulation protocol presented. Therefore, the goal is to find stimulation protocols that maximize perceptual quality. As a proof of concept, we conducted a clinical test on a blind patient implanted with an intracortical 96-channel microelectrode array in their visual cortex. Given the iterative nature of BO, stimulation protocols were presented to the patient iteratively, upon which the patient provided quantitative feedback for the quality of the phosphene perception raised based on a Likert scale, where a score of 7 indicated the highest quality and 0 no perception. To evaluate BO’s performance against a benchmark, another experiment where stimulation protocols were randomly generated was conducted.

In order to apply BO in the context of stimulation protocols, we parametrized the problem space by electrode location indices where a current value was assigned for each indice that was later mapped to a location in the electrode array based on a predetermined list of electrodes. Hence, at each iteration, a current value between [0-50] *µ*A was allocated for each electrode, without surpassing the maximum total current constraint of 500 *µ*A. Whereas standard BO methods do not scale well to problems with over a dozen parameters, here we utilized high-dimensional BO with trust regions to optimize for 40 parameters, thus allocating current only to a predetermined set of 40 electrodes in the array.

Overall, our study provides two unique contributions by proposing an optimization procedure of neural stimulation via cortical neuroprostheses for the blind based on direct patient feedback informing on perceptual quality, and via the use of Bayesian optimization that can scale well to a higher dimension of parameters than the standard BO, through the use of trust regions.

## 2 Methods

### 2.1 Bayesian optimization of patient-specific stimulation for phosphene vision

Here, we propose an experimental design to find optimal patient-specific stimulation parameters for cortical neuroprosthetic vision using high-dimensional Bayesian optimization with trust regions. The design of the experimental process for Bayesian optimization of phosphene vision (BOPhos) (Fig. 1) starts with a vector format of the stimulation parameters to be generated by the Bayesian optimization, with each element corresponding to a stimulation current between [0-50] *µ*A for an electrode, subject to a maximum total current constraint of 500 *µ*A for the total of 40 electrodes. This makes BO operate in the high-dimensional space of 40 parameters, corresponding to stimulation currents for each electrode. The stimulation parameters are later mapped to the electrode array based on the predetermined location indices for the electrodes. The generated stimulation protocol is administered through the implant to the patient’s visual cortex, upon which the patient is asked to provide a score between [0-7] for its perception, where 0 indicates no perception, and 7 corresponds to the highest perceptual quality on a Likert scale. This patient feedback is used via BO to update a surrogate model of the patient response, which in turn helps generate a new stimulation protocol at each iteration via an acquisition function to search the parameter space intelligently by balancing exploration and exploitation. Over time, as the surrogate model learns to capture the patient response effectively, it converges on generating more optimal stimulation protocols that maximize patient response. To analyze its performance, the automated and intelligent search process via BO is compared against a random search process as a benchmark.

**Figure 1:**
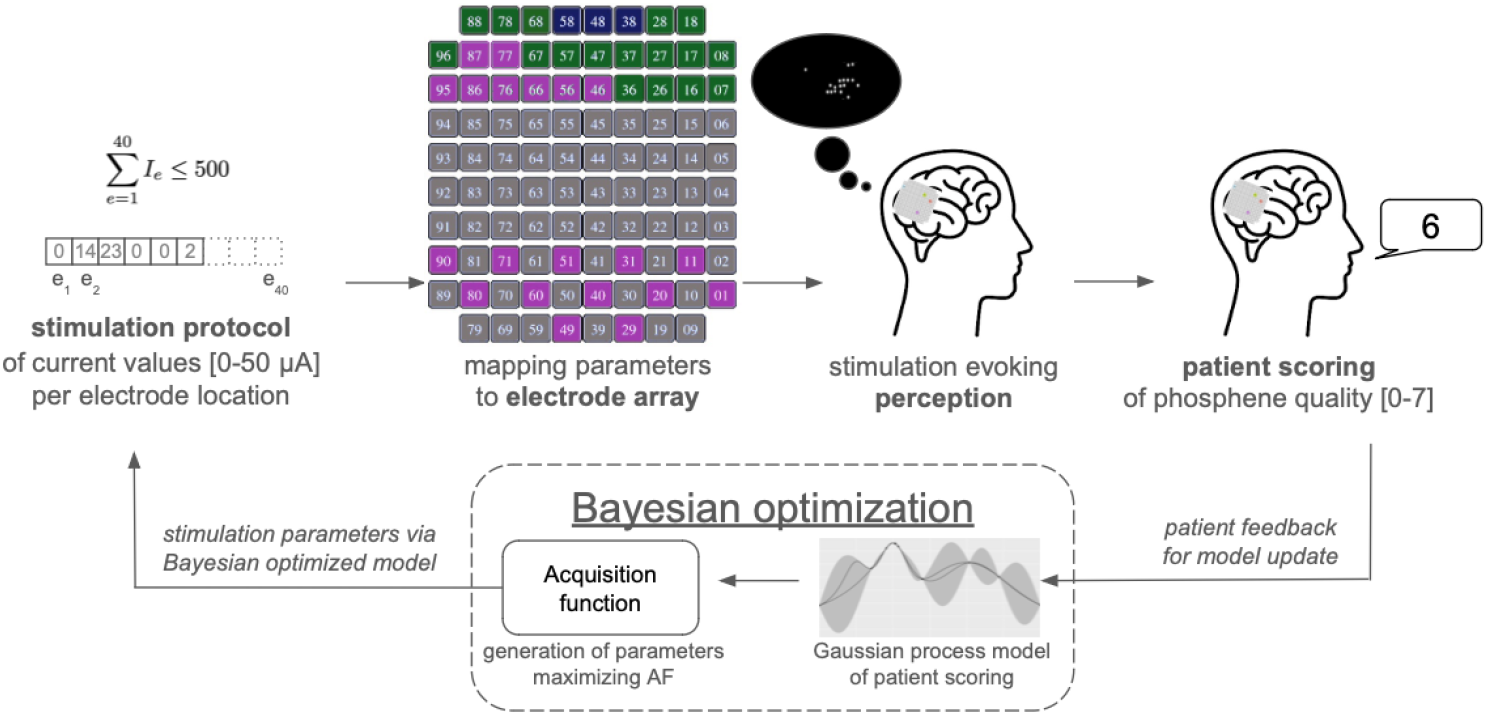
Experimental process of BOPhos involving stimulation protocol generation and stimulation, followed by patient scoring, which can be used as feedback for updating the Bayesian optimization procedure as opposed to the benchmark experiment of random generation of stimulation parameters.

### 2.2 Bayesian optimization

Bayesian optimization (BO) is a framework for solving black-box optimization problems using as few function evaluations as possible (Shahriari et al., 2016). When evaluating the objective function is expensive - as is the case with cortical implant stimulation patterns - it’s crucial to carefully choose where to query next. Efficiently optimizing the objective requires effectively balancing between exploration and exploitation.

Like other forms of optimization, the goal is to find the optimum of a function *f* (**x**) on a bounded set Ω ⊂ ℝ^*d*^. What sets the Bayesian approach apart is the use of a global surrogate model of the objective function, that captures both the predicted objective value and their associated uncertainties. Deciding where to query the objective function next involves a trade-off between exploiting the best function value observed so far and exploring new inputs that might yield even better results. This balance is captured by the acquisition function *α*(**x**; 𝒟_*n*_), which determines the next optimal point by explicitly weighing exploration against exploitation. Instead of using a local gradient or Hessian approximation, the acquisition function leverages *all* the information available from previous iterations when selecting a new input. Together, the probabilistic surrogate model and the acquisition function form the two key components of the Bayesian optimization framework, enabling the optimization of black-box objective functions with a minimal number of evaluations.

#### 2.2.1 Gaussian process surrogate model

The Gaussian process (GP) is a prior probability distribution over functions commonly used as a surrogate model in BO due to its data efficiency and ability to quantify uncertainty. GP regression can predict a function value *f* (**x**) and its uncertainty at an unknown location using a set of past observations 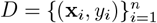.

Assuming noisy measurements, the mean and variance of the prediction at a new input **x** are given by:

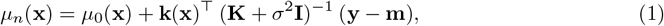

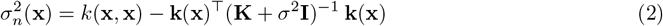

where *µ*_0_(**x**) is the prior mean function, and **k**(**x**) = [*k*(**x, x**_1_), …, *k*(**x, x**_*n*_)] is the covariance vector between the new point and the observed data points. The elements of the prior mean vector and *n × n* covariance matrix are defined as *m*_*i*_ := *µ*_0_(**x**_*i*_) and *K*_*i,j*_ := *k*(**x**_*i*_, **x**_*j*_) respectively.

The posterior mean *µ*_*n*_(**x**) and variance 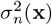 evaluated at a point **x** represent the model’s prediction and its uncertainty after *n* observations respectively. The latter is particularly valuable in BO, as the global optimization approach leverages the uncertainty estimates to balance exploration and exploitation effectively.

The covariance function *k* determines the structural properties of the function modeled by the GP, such as smoothness and periodicity. It acts as a prior over function space, determining how the function behaves between observed data points (Williams and Rasmussen, 2006). Common choices for covariance functions include the Squared Exponential (SE) kernel and the Matèrn 5/2 kernel:

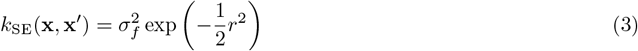

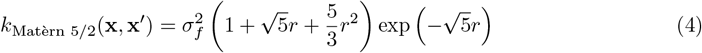

where 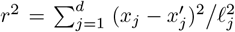 and 𝓁_*j*_ represent the kernel length scales that act as smoothness parameters, and *d* is the dimensionality of the input.

#### 2.2.2 Acquisition function and input constraints

The acquisition function in BO determines where to query the objective function next. Various acquisition functions have been proposed in the literature, each effective under different conditions (Brochu et al., 2010b). Here, we focus on Thompson sampling (Thompson, 1933):

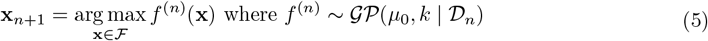

with *f* ^(*n*)^ the sampled function value from the GP after *n* iterations. To select the next input point **x**_*n*+1_, *r* candidate points are sampled. For each candidate point, a function value *f* ^(*n*)^(**x**_*i*_) is drawn from the model’s posterior, where 1 ≤ *i* ≤ *r*. The next input query **x**_*n*+1_ is then chosen as the candidate with the highest sampled function value. This approach naturally balances exploration and exploitation by favoring input points that are both promising and uncertain. This increases the chances of exploring areas with potential for improvement while reducing focus on less promising regions.

##### Generation of candidate points within input constraints

To ensure patient safety, the total current from all stimulated electrodes must remain below a chosen threshold. This allowable maximum total current is often considerably lower than the possible maximum, leading to a trade-off between applying a high current to a single electrode or distributing lower currents across multiple electrodes. We express this as a linear inequality constraint:

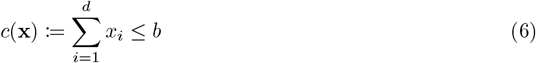

where *d* is the number of electrodes and *b* is the maximum total current threshold. The set of all electrode configurations that satisfy this constraint is given by ℱ = {*x* ∈ Ω | *c*(**x**)}. In our experiment, *d* = 40 and *b* = 500 *µ*A.

To confine the Thompson sampling method to this feasible set, we apply rejection sampling to generate candidate solutions. By sampling *r* candidate points **x**_*i*_ from the full compact set Ω ⊂ ℝ^*d*^ and discarding those that fall outside ℱ, we obtain a set of candidate solutions within the feasible region.

### 2.3 Trust region Bayesian optimization for high dimensional search space

Traditionally, BO has been applied to problems of relatively low dimensionality, typically involving up to a dozen input parameters. While BO has demonstrated highly competitive performance on such low-dimensional search spaces, its application to black-box functions with high-dimensional search spaces remains challenging (Binois and Wycoff, 2022). This difficulty arises due to the exponential growth of the search space with increasing dimensionality and the tendency for functions to be heterogeneous, making it more difficult to fit an accurate global objective function.

To address these challenges, trust region Bayesian Optimization (TurBO) combines principles of local optimization with the global exploration capabilities of BO (Eriksson et al., 2019). This method restricts the search space to a confined region called the trust region, which dynamically moves around the objective landscape. The trust region is centered around the best evaluation found thus far and adapts dynamically by expanding or shrinking depending on whether successive observations are improving or declining, respectively. Typically, the trust region is chosen as a hypercube, although recent work has explored alternatives, such as using a truncated normal distribution to generate candidate solutions (Rashidi et al., 2024).

In this work, we leverage the trust region approach to scale the optimization to a problem with *d* = 40 input parameters representing the electrode currents, which is significantly more than the number of input dimensions that can effectively be handled by traditional BO. Due to the limited sample budget, we utilize a single trust region.

### 2.4 Optimization procedure via BOPhos

The algorithm can be summarized as follows:

1. Evaluate a set of *n*_init_ initial stimulation parameters using space-filling design such as a Sobol sequence, and initialize the trust region at the configuration with the highest patient rating.
2. Repeat until the number of patient evaluations is exhausted:
3. Fit a Gaussian Process model to the observed patient ratings.
  a. Generate a set of *r* candidate points *x*_1_, …, *x*_*r*_ ∈ Ω within the trust region, and discard the samples that fall outside the feasible set (see Equation 6).
  b. For each remaining candidate, sample a realization *f* (**x**_*i*_) to determine the electrode configuration that produces the highest sampled function value.
  c. Present the stimulation protocol and receive the patient’s perception quality rating.
  d. Move the center of the trust region to the highest evaluation and adjust its size based on the success and failure counters.

### 2.5 Experimental setup

#### Subject

The patient tested was a 27-years-old male, who was participating in a 6-month clinical study aimed at developing a cortical visual neuroprosthesis for the blind. The patient was blinded as a result of a traumatic head injury and underwent bilateral enucleation of the eyes one year prior to the experiment. The study received approval from the Clinical Research Committee of the General University Hospital of Elche and was registered on ClinicalTrials.gov (NCT02983370). It adhered strictly to all applicable ethical standards, including clinical trial regulations (EU No. 536/2014, which replaced Directive 2001/20/EC), the Declaration of Helsinki, and the European Commission Directives (2005/28/EC and 2003/94/EC). The patient gave his informed written consent prior to participating. The data was systematically and securely stored.

#### Electrode array and implant location

An intracortical 96-channel microelectrode Utah array, supplied by BlackRock Microsystems (Salt Lake City, UT, USA), was implanted via surgery into the patient’s early visual cortex, between V1 and V2. The procedure followed a standard surgical approach, enhanced by robot assistance (Fernandez et al., 2021; Rocca et al., 2023). Figure 1 includes a picture of the electrode array. A predetermined set of 40 electrodes were included in the study for stimulation (represented in non-gray colors). For the BO implementation, the electrode locations were provided in a fixed order for the mapping of location indices to electrodes as follows: [18, 28, 38, 48, 58, 68, 78, 88, 8, 17, 27, 37, 47, 57, 67, 77, 87, 96, 7, 16, 26, 36, 46, 56, 66, 76, 86, 95, 1, 20, 40, 60, 80, 11, 31, 51, 71, 90, 29, 49].

#### Selection of electrodes and their characteristics

The selected electrodes composed two equally sized main electrode categories: those with measurable phosphene thresholds when stimulated in isolation and those without individual phosphene thresholds. The categorization was made based on electrodes’ observed qualities in previous clinical studies aimed at gathering characteristics of perceptions produced based on patient descriptions upon stimulation. How these electrodes behave in multi-electrode stimulation settings is however not fully known, due to the difficulty of its analysis.

The first group of electrodes (*N* = 20) have a known individual phosphene threshold, meaning they are known to produce phosphenes when stimulated individually above the threshold. This group can be further divided into two subgroups. The first subgroup comprises the three most effective electrodes (*N* = 3) due to their significantly lower thresholds, while the second includes the rest of the electrodes in the group (*N* = 17), as shown in Fig. 1 by colors navy and green respectively.

The second group of electrodes (*N* = 20), colored in pink in Fig. 1, do not have individual phosphene thresholds, hence they do not produce phosphenes when stimulated individually. However, they can play a role in producing perception when stimulated along with other electrodes, as they were previously observed to produce a conscious perception when stimulated concurrently.

#### Electrical Stimulation

We stimulated the visual cortex using Ripple Neuromed’s Explorer Summit processor, which had 3 Micro2+Stim front ends with 32 channels each. The stimulation step size was set to 1 *µ*A. The electrical stimulus lasted for a total duration of 167 ms with 50 pulses at a frequency of 300 Hz. The cathodic-first biphasic pulses had a pulse width of 170 *µ*s with 60 *µ*s interphase duration. The current intensity for a stimulated electrode is generated via the experimental protocol of BOPhos. In order to reduce the total charge injected, we interleaved each stimulation pulse from different electrodes with a time delay of 0.5 ms between the onset of each consecutive pulse inside a train. With a frequency of 300 Hz and a pulse duration of 0.4 ms, up to 6 pulses can be sent separately. For stimulation patterns consisting of more than 6 electrodes, some of the pulses were sent at the same time. The consecutiveness was based on the order of the mapped electrode indices, where the sequence of delays are repeated after every 6 electrodes.

#### Hyperparameters and algorithmic details of Bayesian optimization

For the GP model, we use a Matèrn 5/2 kernel with learnable length scale parameters for each dimension, through a procedure known as Automatic Relevance Determination (ARD) (Neal, 2012). The hyperparameters are determined by optimizing the log marginal likelihood, which is given by:

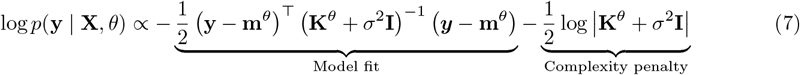

where the trainable parameters *θ* consist of the kernel hyperparameters 𝓁_*i*_, the kernel variance 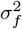 and the (constant) mean function *µ*_*n*_, and 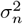 is the noise level that is added to the diagonal of the covariance matrix.

The initial set of *n*_init_ = 12 data points are selected using a Sobol sequence on the full input domain Ω. After observing the patient ratings, the trust region size was initialized at 80% of the input range. At each iteration a set of *r* = 5000 candidate points were generated within the trust region before going for the constraint feasibility check. The experiment was continued until the trust region size reached below 0.045. To adjust the trust region size, a success tolerance of 3 was used (Eriksson et al., 2019).

#### Code implementation

All code was implemented in Python, and is available at https://github.com/burcukoglu/BOPhos.git.

#### Baseline experiment

To compare the performance of Bayesian optimization in converging to the generation of optimal stimulation parameters, a second experiment was conducted as a benchmark with random generation (RG) of stimulation parameters.

## 3 Results

### 3.1 Convergence properties

BO has converged to optimal parameters for generating stimulation protocols that lead to highly rated perceptions in only 230 iterations within 20 minutes of patient testing, demonstrating its efficiency in the search process. For comparison, the RG experiment was also run for 230 iterations.

### 3.2 Patient scoring in response to stimulation protocols

The stimulation protocols generated by Bayesian optimization are compared to those randomly generated across the iterations or trials for each experiment. Patient scoring across trials are displayed in Figure 2, which shows a higher maximum score reached by the BO experiment than the RG experiment. Cumulative scores show that with BO, the patient scoring increases steadily and reach to more than two times the cumulative patient scoring in the RG experiment. The actual patient scoring demonstrates the unsystematic search of the RG approach over the parameter space, which produces lower and inconsistent scores with an initial dip of scores at the beginning. Stimulation protocols that could lead to scores beyond the mean of the possible score range are barely generated, suggesting that only suboptimal stimulation protocols had the opportunity to be tested for patient perception. In contrast, BO yields a steady increase to reach higher scores soon after the start of the experiment, and maintains the high scores with consistency across many iterations, yet with regular drops to its value. The patient scoring demonstrates that BO is effective in generating stimulation protocols whose perception quality is evaluated at the highest levels by the patient. However, due to its nature as an iterative optimization procedure, it is prone to effects of adaptation due to repeated stimulation, as suggested by the regular patient scoring drops across later iterations despite the overall consistency in consecutive scores.

**Figure 2:**
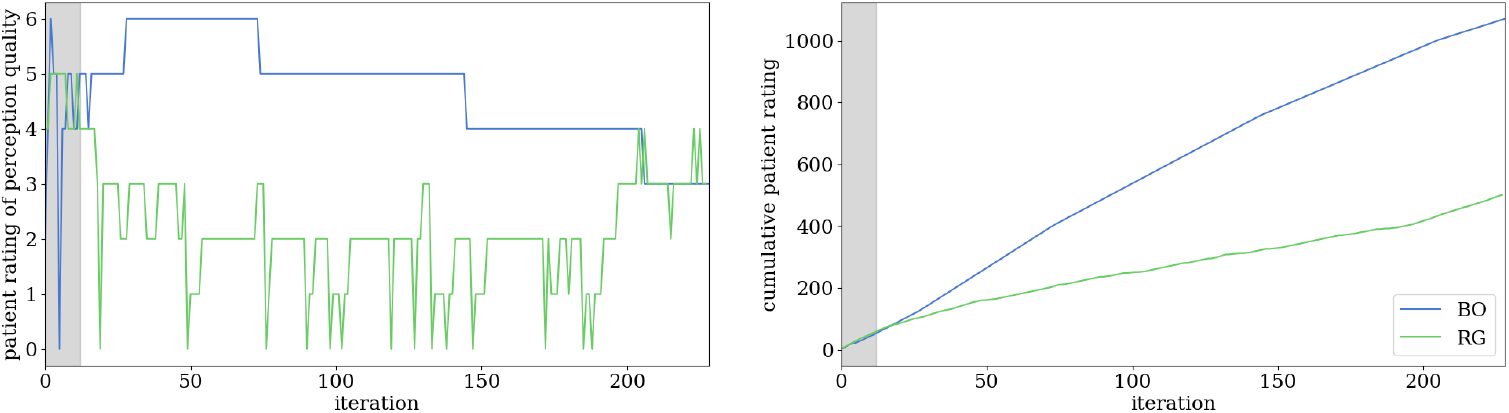
Patient scoring across iterations of stimulation protocol generation (left), and cumulative patient scoring (right). The gray area indicates the initial set of data points tested for the BO experiment, before the optimization begins.

### 3.3 General characteristics of the stimulation protocols

Total currents across trials for the two experiments are compared in Figure 3. In the RG experiment, the stimulation protocols generated produce inconsistent total current values, yet reaches the maximum total current constraint of 500 *µ*A a few times. BO, however, converges to the maximum total current soon after the iterations start. While high levels of current are not necessarily desirable for repeated stimulation due to the risk of overstimulation, it was observed in clinical studies that patients have a preference for brighter stimuli, which is attributed to higher stimulation currents. This explains why the BO converged on stimulation protocols with the highest total current possible, within the safety limits provided for it. It is worth noting that in the RG experiment, despite receiving the highest total current a few times throughout the iterations, the patient has not evaluated these stimulation protocols with higher scores. This emphasizes the importance of the choice of electrodes in reaching optimal stimulation protocols, which the BO has clearly succeeded in, given the difference in patient scoring to the stimulation protocols with the highest total currents compared to the RG experiment. As electrodes could have different phosphene perception thresholds, beyond which a stimulation could produce a perception only, it could be the case that despite high total currents administered, some of it has not contributed directly to patient perception, hence causing a lower patient scoring. However, BO seems to overcome this by generating stimulation protocols of not only high total current but also with a choice of electrodes that have low phosphene thresholds, hence more effective in producing perception. This is later further analyzed based on the knowledge of individual perception thresholds for electrodes acquired in previous clinical studies.

**Figure 3:**
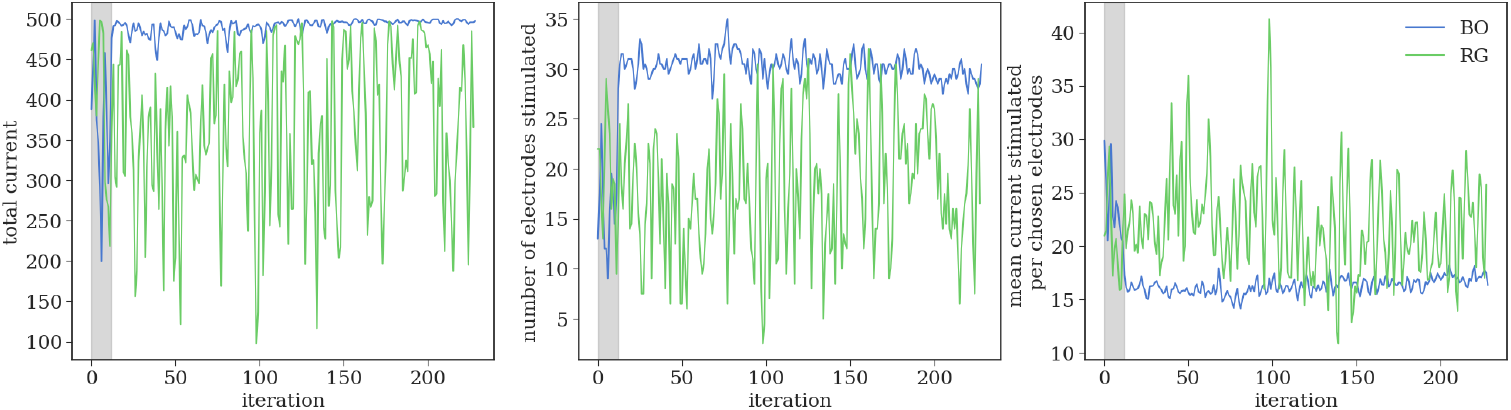
Total current across iterations (left), number of electrodes stimulated per generated stimulation protocols (middle), and mean current per stimulated electrode (right). Plotted values show the moving average with a window size of 2. The gray area indicates the initial set of data points tested for the BO experiment, before the optimization begins.

Looking at Figure 3, we see that the number of electrodes stimulated is also inconsistent for the RG experiment, whereas it converges to around 30 for the BO out of 40 possible electrode choices, soon after initial iterations. The mean current per stimulated electrodes in a generated protocol is similarly inconsistent for the RG experiment, yet converges to around 16 *µ*A for BO. While the number of electrodes stimulated is in general higher for BO, the mean current per stimulated electrodes is also generally lower, despite higher total current values on average. This may suggest an advantage of the BO framework for minimizing the mean current per electrode stimulated, despite not being explicitly steered to optimize for it.

### 3.4 Analysis on electrode categories

The choice of electrodes by the generated stimulation protocols and their currents are further investigated based on the categorization of electrodes. As Figure 4 shows, the choice of the most effective subcategory of electrodes and also their current allocation is much higher in the BO compared to the RG experiment, demonstrating the success of BO in learning to generate the most optimal electrodes for phosphene perception. Similarly, the selection and the current allocation for the main category of electrodes with individual phosphene perception thresholds were significantly higher in the BO than in the RG experiment. Within this main category, BO especially learns to allocate most of the current it distributes to the subcategory of most effective electrodes, to the extent that the second subcategory is allocated less current in BO than in RG, yet still with an increased count of stimulated electrodes. This demonstrates BO’s effectiveness not only in the selection of more effective electrodes but also in their allocation of higher current values. On the other hand, the electrodes without individual phosphene thresholds seem to be selected more in BO, and also with higher current allocation than any other group, despite being selected less by BO compared to the other main category that contains equal number of electrodes.

**Figure 4:**
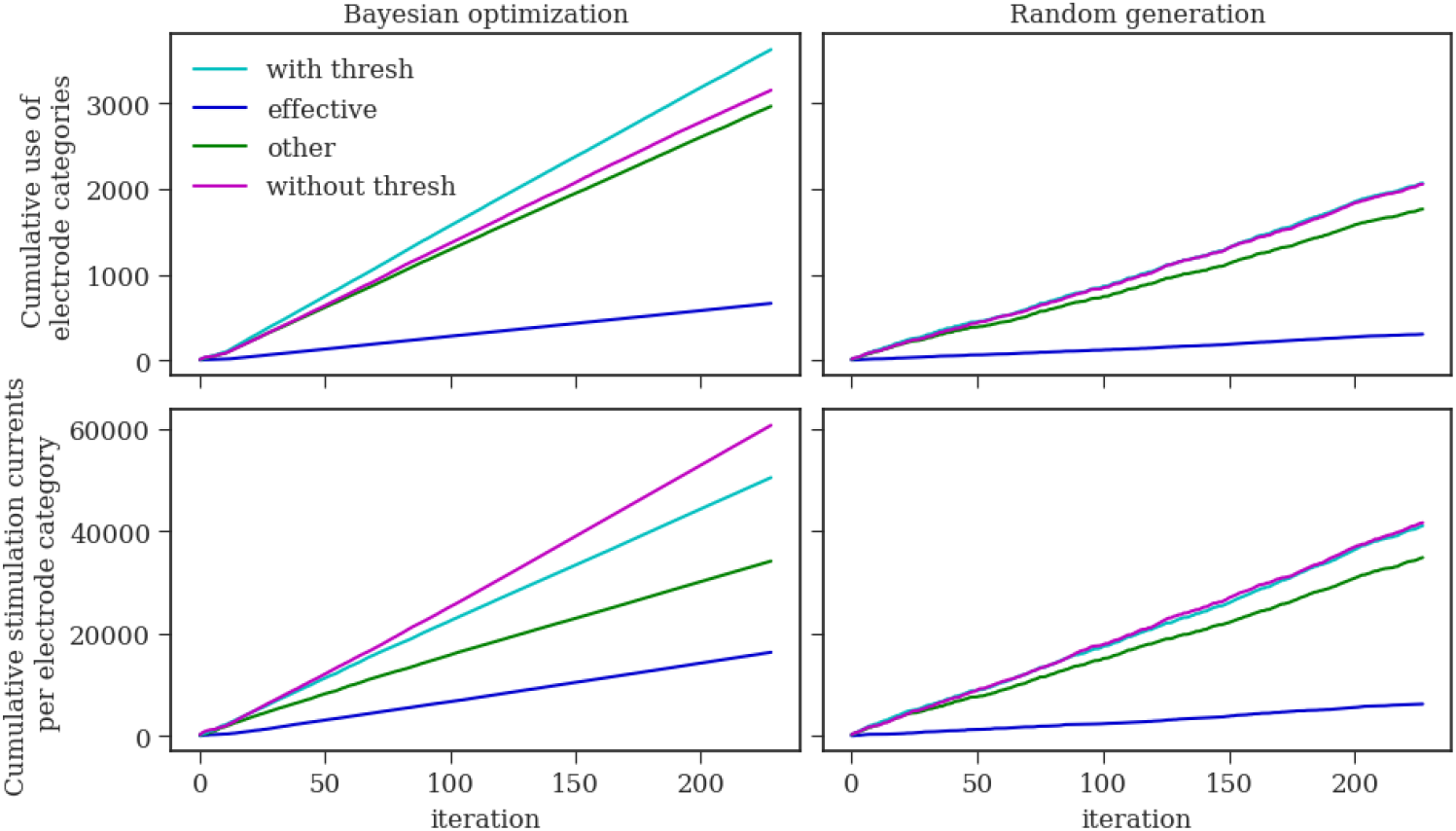
Cumulative count of electrodes stimulated per electrode category (top) and their cumulative current (bottom). The two main categories are electrodes with individual phosphene thresholds (*with thresh*), and without individual phosphene thresholds (*without thresh*). The first category (*with thresh*) is further divided into two subcategories, with the first (*effective*) including the most effective electrodes due to lower individual phosphene thresholds, and the second (*other*) including the rest of the electrodes in the category. Note that the number of electrodes across categories differ.

While this preference for electrodes without individual phosphene thresholds in current allocation may reflect an important contribution to perception from them in multi-electrode stimulation settings, it can’t be concluded confidently due to a lack of an approach of clear analysis. As an alternative explanation, it could be the case that for multi-electrode stimulation settings, these electrodes have high phosphene perception thresholds, hence making the stimulation current really high, yet with minimal contribution to the brightness or perception. While the patient scoring seems to increase initially, it is observed to decrease later, which might be a result of higher allocation of currents to electrodes without individual phosphene thresholds, in the case their stimulation is ineffective for perception. However, the decline in the patient scores for later iterations of BO could also very well be due to adaptation effects or psychological aspects like motivational and attentional drop in the patient especially due to the regular, gradual nature of the decline that rather maintains the consistency of scores in between. Ideally, future analysis should support the current observations with an analysis of the impact of adaptation and psychological effects, and also the testing of BO-generated stimulation protocols in the later iterations before any exposure to prior stimulation. In order to get more information about the choice of electrodes, Figure 5 informs us about the non-cumulative scoring of the electrode category count and currents per category. RG experiment reflects the randomness in the generation of stimulation protocols, where electrode selection and current allocation per category is inconsistent across iterations and merely reflect the number of electrodes available in each category in their ranking of selection count and total current. In contrast, it can be observed that the three most effective electrodes were always chosen by BO from right after the start, with the highest mean current per electrode overall across categories. This shows BO’s success in selecting and allocating higher current to the electrodes with the lower individual phosphene thresholds. The count of electrodes stimulated for the rest of the categories rather reflects the size of the categories, with on average 16 out of 20, 13 out of 17, and 14 out of 20 electrodes being chosen for remaining groups, respectively for the main group with threshold, the subgroup of those with threshold, and finally those without threshold. Still, the group of electrodes without individual thresholds takes the attention, with the group’s total current surpassing those of the group of electrodes with individual thresholds, despite the lower selection ratio for the electrodes in the former. As a result, the group of electrodes without individual thresholds reaches to the second highest levels of mean current allocation per electrode, following after the group of most effective electrodes.

**Figure 5:**
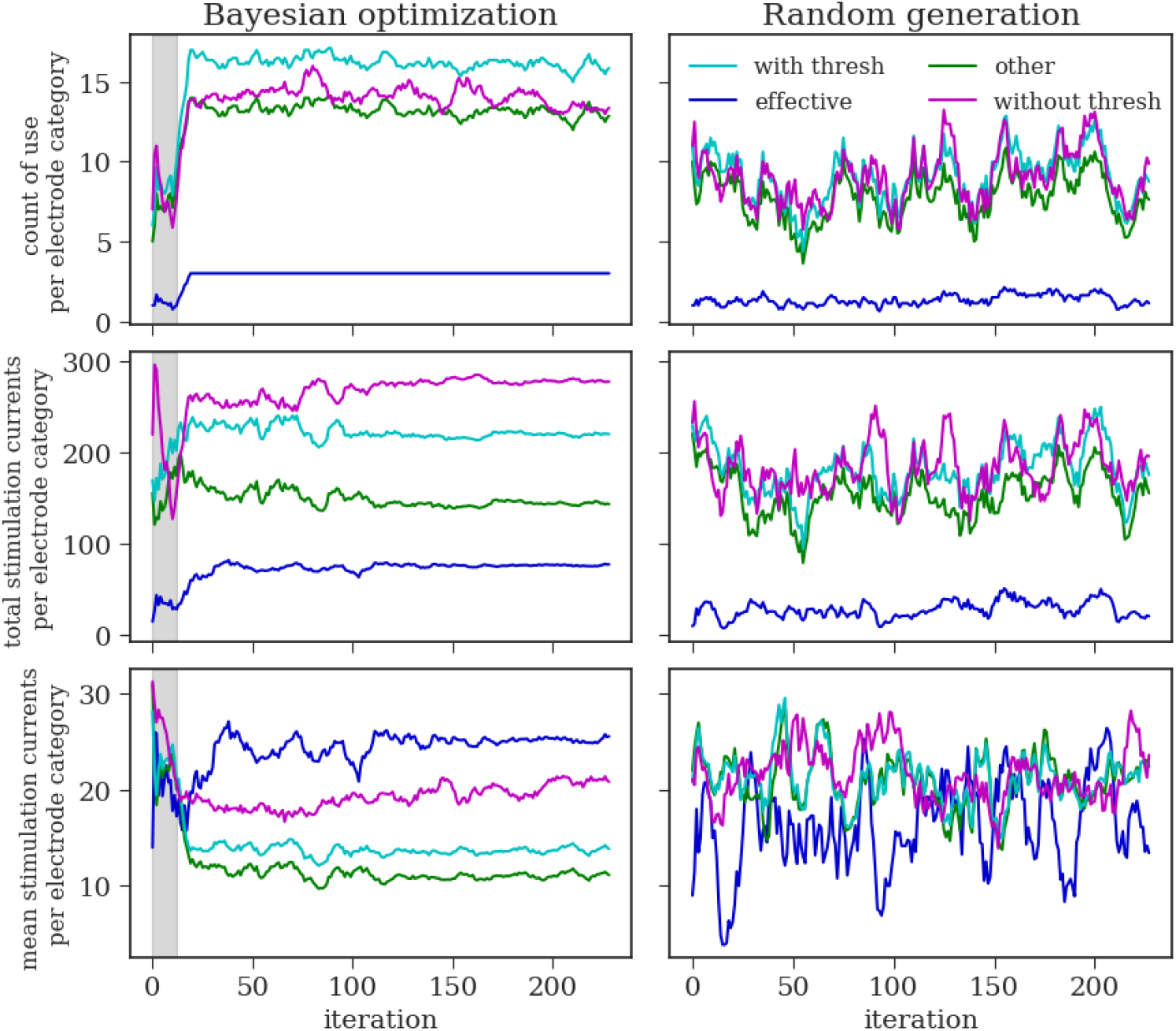
Number of electrodes stimulated per electrode category (top), their total currents (middle) and the mean current per electrode category (bottom). Number of electrodes available in the two main categories, and later for the two subcategories of the first group are as follows: *with thresh*=20, *without thresh*=20, *effective*=3, *other*=17. Plotted values show the moving average with a window size of 8. The gray area indicates the initial set of data points tested for the BO experiment, before the optimization begins.

### 3.5 Analysis on individual electrodes

An individual analysis for each electrode within each category is provided in Figure 6. While each electrode is selected by and allocated current to roughly at the same levels for the RG experiment, a clear preference for or aversion to certain electrodes is apparent in the BO experiment, reflecting their contribution levels to perception or phosphene thresholds. Among the most effective electrodes, all are chosen roughly equally, with electrode 38 being allocated the most current by BO and electrode 48 the least. As for their stance in the bigger group of electrodes, there are 8 other electrodes in the with thresholds category, and 12 in the without thresholds category, that are selected to be stimulated as often. Among the electrodes with individual thresholds, the three most effective electrodes are all in the top six (along with 8, 17 and 78) of the most current-allocated ones, with 38 having the highest current allocation of the main category. Surprisingly, 2 other electrodes (60, 51) have higher current allocation than 38, which are part of the category of electrodes without individual thresholds, implying a high contribution in perception on their part in the multi-electrode settings.

**Figure 6:**
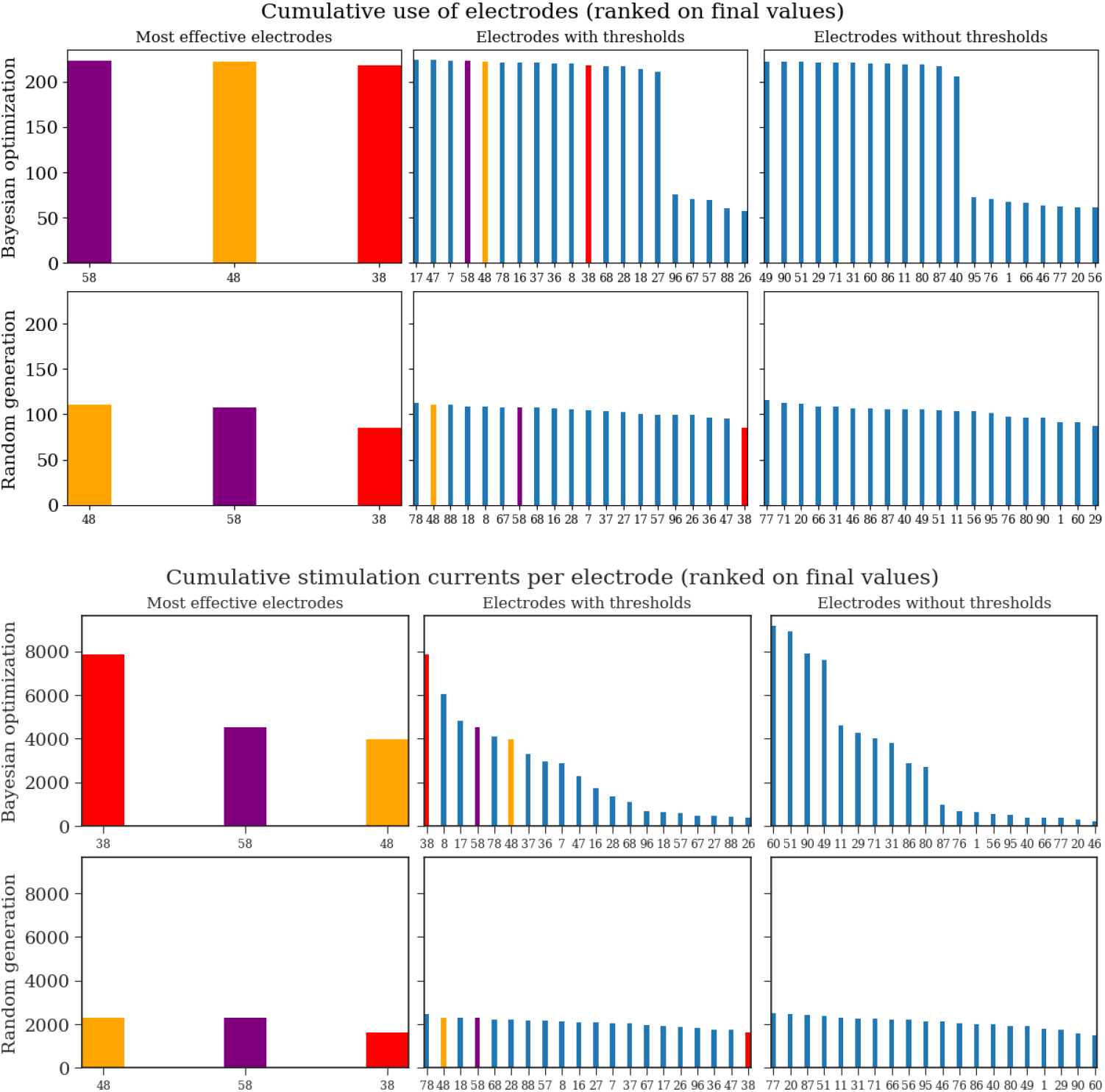
Cumulative count of stimulation per electrode (top) and their final cumulative current (bottom) grouped based on electrode categories. Electrodes are ranked by their value for each subplot from the highest to the lowest.

There is a clear aversion of BO to certain electrodes, which have been allocated very little current across iterations and only infrequently selected. These electrodes seem to have minimal contribution to perception and thus can be considered rather ineffective. Among the electrodes without individual thresholds, there are more electrodes BO is averse to (95, 76, 1, 66, 46, 77, 56, 20) based on both current allocation and electrode selection, compared to the with thresholds category. The electrodes with individual thresholds show the most dispersed current allocation levels, with half of them being allocated rather low current levels, and five having both the lowest current and selection levels (26, 88, 67, 57, 96). A few electrodes in these two categories stand out for having both the lowest current and yet highest selection levels. These are electrodes 40 for the first, 27 and 18 for the second category respectively, in the order they were just mentioned. It may imply rather a consistent contribution to perception by these electrodes in the multi-electrode settings, even with low current stimulation. Overall, BO seems to provide rather a fast way of gathering insights on specific electrodes’ contribution to perception. Regarding the previously discussed results, the higher current allocation levels for the top ranking electrodes without thresholds, as opposed to higher number of effective electrodes in the with individual thresholds category, could be the reason for the former category being allocated higher total and mean current than the latter category.

In conclusion, aligned with the findings of previous clinical studies, BO learnt to converge on choosing and allocating the most current to the most effective electrodes. Furthermore, it enabled a way to test for optimal stimulation protocols in the multi-electrode setting in a fast and efficient manner, which would have required an exhaustive search if it was to be done purely via clinical studies. This could make the onboarding of a patient newly implanted with a cortical neural prosthesis faster, allowing for efficient and systematic initial experimentation to reach conclusions about the effectiveness of electrodes and their phosphene perception thresholds in a patient-specific manner. Moreover, going beyond merely quantifying perception thresholds and assuming sufficient perception with currents above, BO helps inform about the quality of above-threshold perception, which can be good or poor irrespective of the level of current administered even in the case of a low threshold, thus bringing additional value to the current capabilities of clinical studies.

## 4 Discussion

This study proposed BOPhos, an experimental procedure to optimize stimulation protocols for cortical neuroprosthetic vision based on direct patient feedback on perceptual quality using Bayesian optimization, in order to efficiently and systematically search through the huge parameter space of multi-electrode stimulation in safety-critical clinical settings. By converging within as short as 20 minutes, BO has learned to generate optimal stimulation protocols that lead to highly rated patient perceptions, through determining which of the 40 electrodes to stimulate and with how much current, under safety constraints. This proof-of-concept study demonstrates the power of high-dimensional BO with trust regions in quickly converging to optimal stimulation protocols based on direct patient feedback, providing a more efficient search for stimulation parameters in clinical settings. Compared to random search, BO has learnt to allocate less current per stimulated electrode, despite higher total current allocation and a higher number of electrodes stimulated on average, whereas it has learnt to allocate higher current for more effective electrodes. Thus, BO has been shown to be useful, especially for identifying effective electrodes and informing about perception quality in multielectrode stimulation for fast onboarding of patients in clinical settings, given its alignment with the findings of the previous clinical studies. Therefore, BO provides an efficient method for initial systematic experimentation with a patient for testing the effectiveness of multi-electrode stimulation protocols, which could have otherwise been an exhaustive and time-consuming search process.

While the impact and benefits of the BO are clear in this proof-of-concept study, due to the iterative nature of the algorithm, the patient is required to be stimulated repetitively. While this repetition is aligned with the inherent design of the cortical visual neuroprosthesis, it could be a challenge in interpreting the performance of the BO framework in the context of neural stimulation, where model updates are based on patient feedback. Due to adaptation effects as a result of repetitive stimulation or psychological aspects like motivational and attentional drop, patient ratings of the stimulus may reduce over time stemming from a decrease in the responsiveness of the sensory system or in the interest of the patient to continue the experiment. While these effects may make it harder for BO to get accurate feedback on patient perception, thus slow down its convergence or cause it to converge to a suboptimal model of the patient response, it still manages to converge towards the right direction due to an overall decline in patient ratings. Ideally, however, such adaptation effects should be taken into account by adjusting the stimulation over time so that sufficient current is administered to ensure the perception of the same stimuli with a higher current over time. Alternatively, a followup study could be done on electrodes found to be more effective based on BO, in order to test their contribution to perception more accurately, based on a selection from a smaller subset of electrodes and before any exposure to prior stimulation. Another means to deal with adaptation and psychological effects could be by administering the iterative stimulations with a greater time interval in between, or with regular breaks between multiple consecutive stimulations, to minimize the side effects of repeated stimulation.

In order to reach more definitive conclusions about the effectiveness of electrodes and current allocation required for highly rated perception, some limitations should be taken into account and mitigated for future work, apart from the adaptation and psychological effects. First of all, due to the limited availability of the implanted blind patients, the study was conducted only with a single patient; hence, whether the use of BO can generalize to more patients could not be investigated. Second, the time of testing available with this single patient was also limited, which allowed us to only run both experiments once, with BO followed by the RG experiment immediately after. Ideally, for each experiment, the same experimental design could be repeated multiple times with enough time difference in between so that more conclusive results could be reached about the effectiveness of electrodes and current allocation required for highly rated perception over multiple runs. Even if this were not possible, an alternative would be having both experiments again but this time with RG being tested before BO to eliminate the effects of order. However, limited patient testing time has not made these ideal scenarios possible, thus potentially causing a bias in our results more in favor of the BO experiment due to less exposure to prior stimulation, hence possibly leading to higher patient ratings or more attentiveness to perception.

Apart from these limitations, another means to reach more definitive conclusions could be through incorporating information on individual phosphene thresholds for the electrodes, so that it can be known roughly how much of the current allocated per electrode is effective, especially for electrodes that always have individual phosphene thresholds when stimulated alone. This could be interesting to investigate for future work.

In terms of the design of the Bayesian optimization procedure, different alternatives were considered and could provide a route for future work. First, preferential Bayesian optimization (Brochu et al., 2010a; González et al., 2017), where the patient is asked to provide a preference between two different stimuli at each iteration, can be used for incorporating feedback from the patient, as it is easier for human subjects to judge a relative comparison between two stimuli more accurately rather than judging one single stimulus across a series of stimuli in absolute terms (Miller, 1956; Kahneman and Tversky, 1979; Seymour and McClure, 2008). While in the context of neural stimulation such preferential comparison will still give rise to order effects between the two stimuli, they could be potentially mitigated by the above-mentioned solutions. Second, instead of only maximizing patient perception quality based on patient feedback, a second objective can be incorporated to minimize total stimulation current, since minimizing stimulation current is both desirable for safety-critical settings like neural stimulation and can also compensate for the tendency of BO to converge to maximum total current possible when only optimizing for patient perception. A challenge in combining two separate objectives in this case comes from the patient rating being an ordinal variable, whereas the total current is a continuous variable. Whereas there are studies investigating BO for mixed variable multi-objective problems (Sheikh and Marcus, 2022), an implementation of this in neural stimulation context needs further analysis. Future work can consider these alternatives to potentially improve the effectiveness of optimization pipelines for neural stimulation.

To our knowledge, this is the first study in the context of cortical neuroprosthetic vision that optimizes stimulation parameters *in vivo* directly based on patient feedback, by focusing on the informative value of perception quality for clinical studies. It is also special in optimizing for more than a dozen parameters in the context of neural stimulation, beyond which typical BO tends to fail. We successfully demonstrate a high-dimensional optimization of 40 parameters, by adopting trust region-based BO. It is worth noting that while TurBO provides efficiency benefits in searching through the high-dimensional parameter space, it does so by letting go of global exploration via the use of trust region only around the best evaluation so far. Hence, more optimal solutions might exist, albeit at the cost of efficiency in the search, especially when considering the full array of 96 electrodes as the parameter search space as an obvious next step.

Bayesian optimization can also bring the possibility for closed-loop control of cortical neuroprosthetic vision one step closer through a potential incorporation of neural responses as the feedback to update the surrogate model. Future work should also focus on the incorporation of a visual processing pipeline, where a camera image is processed to generate stimulation parameters for a targeted phosphene vision, to evoke the desired perceptions in the brain of the blind patient for a complete optimization of cortical neuroprosthetic vision via BO. This proof-of-concept study here lays the groundwork for any future work in this direction for optimizing neuroprosthetic vision based on direct patient feedback in a high-dimensional search space.

## 5 Conclusion

We have proposed BOPhos, an experimental design for finding optimal patient-specific stimulation parameters for cortical neuroprosthetic vision using high-dimensional Bayesian optimization with trust regions. We demonstrated the power of BO in quickly converging to optimal parameters *in vivo* by testing on a blind patient implanted with a Utah array in the early visual cortex and using direct patient feedback on perception quality for steering the optimization towards maximizing a perceptual score under safety constraints. Bayesian optimization proved useful, especially in allocating higher current to more effective electrodes and leading to higher perceptual quality ratings. While a complete Bayesian optimization of phosphene vision would also require a visual processing pipeline to evoke targeted perceptions, this study establishes a foundation for future research by incorporating direct patient feedback for phosphene optimization in a high-dimensional search space for the first time. Our study contributes towards an efficient and systematic search for the huge parameter space of multi-electrode stimulation for fast onboarding of patients in clinical settings via an initial experimentation on the effectiveness of electrodes and optimality of stimulation parameters for perceptual quality.

## Acknowledgements

This work has received funding from the European Union’s Horizon 2020 research and innovation programme under grant agreement No 899287 (project NeuraViPeR). This publication is also part of the project Innovative Neurotechnology for Society (INTENSE) with project number 17619 of the Crossover programme. This work was supported in part by grants PDC2022-133952-100 and PID2022-141606OB-I00 from the Spanish “Ministerio de Ciencia, Innovación y Universidades”, and by grant CIPROM/2023/25 from the Generalitat Valenciana (Spain). This publication is also part of the project Dutch Brain Interface Initiative (DBI2) with project number 024.005.022 of the research programme Gravitation which is (partly) financed by the Dutch Research Council (NWO). This publication is also part of the project ROBUST: Trustworthy AI-based Systems for Sustainable Growth with project number KICH3.L TP.20.006, which is (partly) financed by the Dutch Research Council (NWO), ASMPT, and the Dutch Ministry of Economic Affairs and Climate Policy (EZK) under the program LTP KIC 2020-2023. All content represents the opinion of the authors, which is not necessarily shared or endorsed by their respective employers and/or sponsors.

